# Crowding on DNA modulates SSB protein binding mode kinetics

**DOI:** 10.64898/2026.07.02.736164

**Authors:** Alberto Pérez-Mugía, Bruno Marcos, Juan P.G. Villaluenga, Borja Ibarra, Francisco J. Cao-García

## Abstract

Single-stranded DNA-binding (SSB) proteins play a crucial role in DNA replication by binding to single-stranded DNA (ssDNA) in multiple binding modes, depending on conditions such as salt and protein concentrations. The coverage-dependent effects on the kinetics of these binding modes remain incompletely understood. In particular, the bimodal binding kinetics and the further SSB-ssDNA shortening observed when SSB is removed from the media. Here, we develop a kinetic model extending the Tonks-McGhee-von Hippel framework to incorporate ligand crowding and mode transformations, capturing the inhibition of SSB binding and transitions to higher binding modes as coverage increases. This model quantitatively reproduces experimental binding kinetics and coverage-dependent behaviors observed for human mitochondrial SSB (HmtSSB) and E. coli SSB (EcoSSB). Our findings elucidate the impact of ligand crowding on SSB-ssDNA interactions and provide a generalizable framework for studying multimode ligand binding to polymers, with implications for understanding genome maintenance mechanisms.

**GRAPHICAL ABSTRACT:** 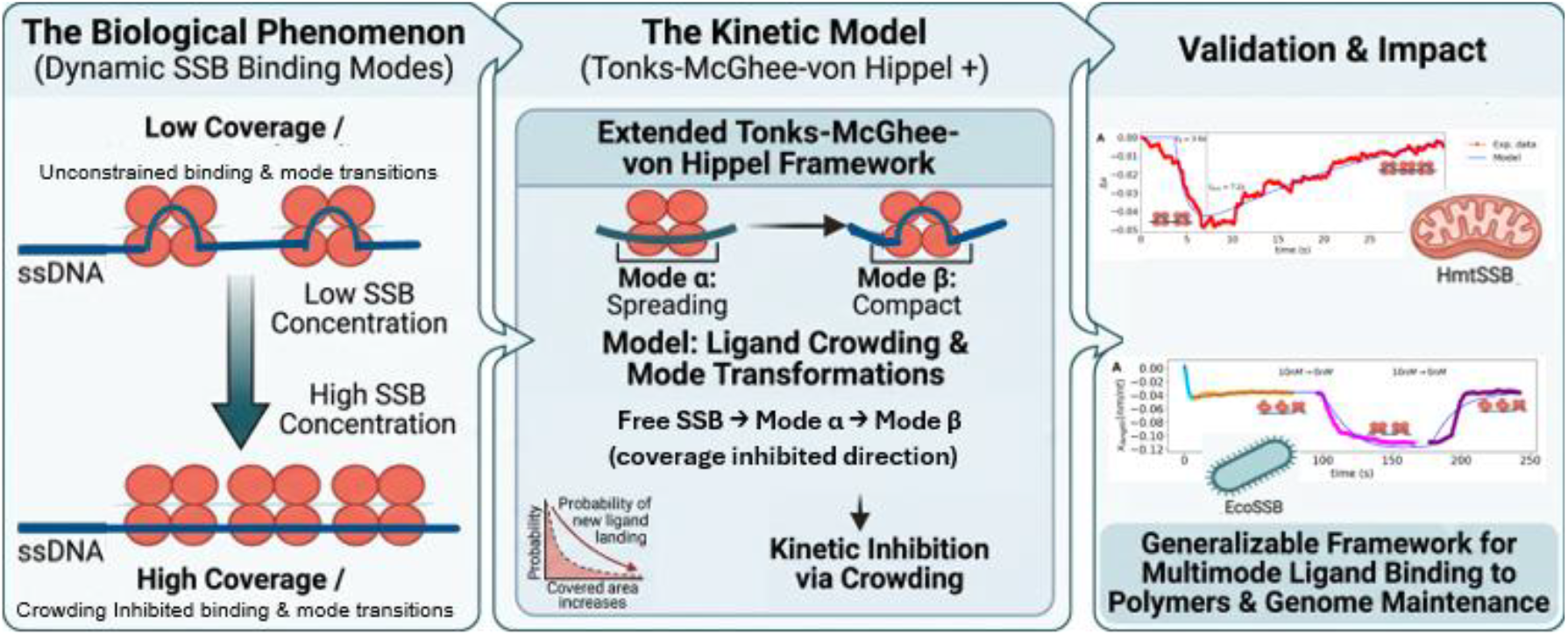

## INTRODUCTION

Single-stranded DNA-binding proteins (SSBs) bind with high affinity and without sequence specificity to single-stranded DNA (ssDNA). SSBs play a crucial role in DNA replication by protecting the lagging DNA strand and preventing secondary structure formation until it can be replicated (Kornberg and Baker 1992). In addition, SSB proteins have been shown to stimulate the activities of various genome maintenance proteins (Shereda et al. 2008), particularly DNA polymerases (Plaza-G.A. et al. 2023).

SSBs can bind to ssDNA in multiple modes, with each mode involving the protein covering a different number of nucleotides (Flynn and Zou 2010; Murzin 1993). The specific binding mode adopted depends on experimental conditions such as salt and SSB concentrations (Antony and Lohman 2018; Ciesielski et al. 2015; Suksombat et al. 2015; Maffeo and Aksimentiev 2017; Morin et al. 2015; Naufer et al. 2021). For example, E. coli SSB (EcoSSB) exhibits several binding modes characterized by varying degrees of DNA wrapping, with transitions between these modes occurring via wrapping and unwrapping pathways (Suksombat et al. 2015; Bonde et al. 2024).

Recent advances using optical tweezers have enabled detailed measurements of the mechanical properties of SSB-ssDNA complexes, providing insights into both the equilibrium coverage (the fraction of ssDNA bound by SSB) and the predominant binding mode under various conditions (Morin et al. 2017; Naufer et al. 2021). Measurements of end-to-end distance and force-extension curves have revealed changes in the structure and dynamics of the SSB-DNA complex (Jarillo et al. 2017).

In Ref. (Morin et al. 2017), the analysis of the experimental data on human mitochondrial SSB (HmtSSB, a close homolog of EcoSSB) binding to ssDNA implied that increasing HmtSSB concentrations in solution leads to higher coverage of the DNA by HmtSSB and induces a shift toward the lower binding mode of HmtSSB. HmtSSB presented two binding modes: a lower mode covering approximately 40 nucleotides and a higher mode covering about 70 nucleotides. The experimental data also revealed a transient shortening of the DNA, likely corresponding to a temporary transition to the higher binding mode.

In Ref. (Naufer et al. 2021), experimental data on EcoSSB binding to ssDNA revealed bimodal binding kinetics, characterized by a transient shortening of the end-to-end extension of the ssDNA-SSB complex, particularly pronounced at low forces. This bimodal behavior is attributed to a temporal transition to a higher binding mode, followed by a return to the lower binding mode. Notably, removing EcoSSB from the solution does not immediately restore naked ssDNA, leading to hysteresis upon reintroducing EcoSSB. These observations are explained as consequences of changes in the effective binding rate and mode transition rates at high DNA coverages (Naufer et al. 2021).

Here, we investigate whether ligand crowding on ssDNA can account for the experimental observations regarding the equilibrium and kinetics of SSB binding (Morin et al. 2017; Naufer et al. 2021), and seek to provide a theoretical framework for the observed changes in effective binding and transition rates as DNA coverage increases.

To address this question, we explicitly include transformations between different SSB binding modes, extending our previous kinetic models of ligand binding (Villaluenga, Vidal, and Cao-García 2020; Villaluenga and Cao-García 2022; Villaluenga, Brunete, and Cao-García 2023), based on the lattice Tonks-McGhee-von Hippel (TGH) equilibrium model (McGhee and von Hippel 1974, 1976).

Here, we show that the resulting complete kinetic model explains the observed SSB-DNA binding kinetics (Morin et al. 2017; Naufer et al. 2021). These results imply that the observed binding kinetics is compatible with crowding-induced inhibition of binding and mode transformation.

## MATERIALS AND METHODS

### Statistical-mechanics lattice models of ligand binding to ssDNA

The polymer is modeled as a one-dimensional array of binding sites, lattice binding model. Our polymer is the ssDNA, and each binding site is a nucleotide. See Fig. 1.

**Figure 1.**
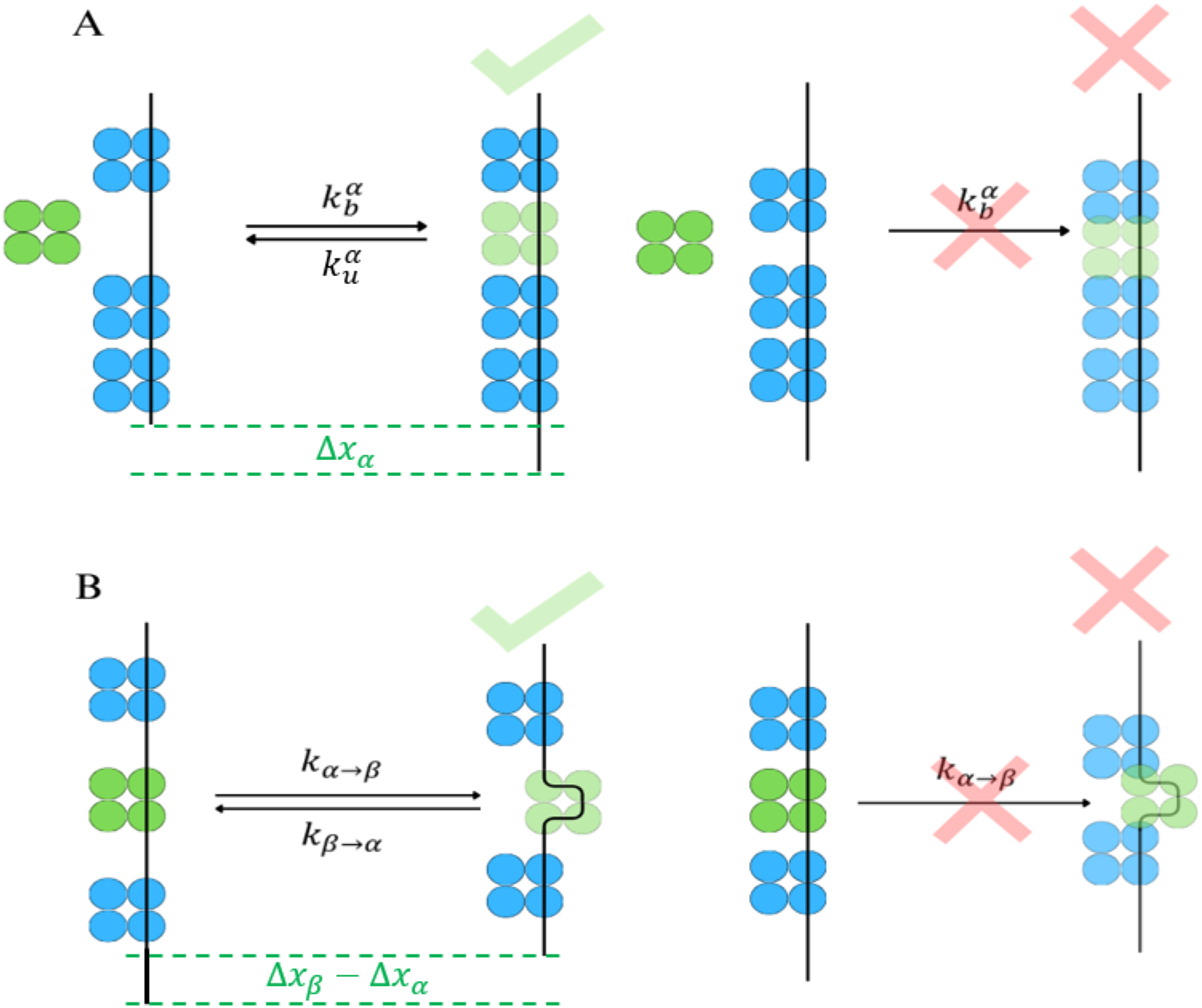
Proposed mechanisms of how ligand crowding in DNA impacts SSB binding kinetics. **(A) Ligand crowding in DNA inhibits SSB binding**. SSB binds to several nucleotides, binding only takes place if enough binding sites are free. As is represented with allowed (left) or forbidden (right) binding of the green ligand. **(B) Ligand crowding in DNA inhibits transitions to higher binding modes**. SSB can bind DNA in different binding modes, higher binding modes are only reachable if the required additional nucleotides are free. As is represented by the allowed (left) or forbidden (right) transition of the green ligand to the higher binding mode. This model leads to saturation-dependent binding and transformation rates, explaining the main features observed in the SSB binding kynetics to DNA. These proposed mechanisms lead to effective inhibited binding and transition rates in the kinetics, which explain the main features of the experimentally observed kinetics (see Fig. 2, 3 and 4)

Equilibrium states are computed in lattice models through different methods. McGhee and von Hippel (McGhee and von Hippel 1974, 1976) proposed a procedure to count the possible binding positions of a large ligand (i.e., one that binds multiple binding sites) on a long polymer (modeled as a one-dimensional lattice). They derived the equilibrium abundances of the ligand in the ligand-polymer complex. Other authors further developed this procedure later (Epstein 1978; Schwarz and Stankowski 1979; Wolfe and Meehan 1992; Lincoln 1998; Tsodikov et al. 2001). Other methods used to determine the equilibrium state of ligand-polymer complexes include the transfer matrix method (Tsuchiya and Szabo 1982; Di Cera and Kong 1996; Nishio et al. 2003; Teif 2007), the generating function method, and the recurrent relation method [see (Teif and Rippe 2010)]

Fewer results are available for the kinetics of irreversible and reversible binding for a single type of ligand (Epstein 1979a, 1979b; Schwarz and Watanabe 1983).

We have previously developed binding kinetic models addressing the binding of large ligands to polymers with phenomenological approaches (Jarillo et al. 2017) and with the McGhee von Hippel counting of binding sites applied to non-cooperative ligands (Villaluenga, Vidal, and Cao-García 2020), and cooperative ligands (Villaluenga and Cao-García 2022). We also considered the case of several competing ligands (Villaluenga, Brunete, and Cao-García 2023).

However, a kinetic model for SSB binding to DNA must account for the transitions between SSB’s different binding modes to DNA. These binding mode transformation terms have not been considered in the previous kinetic studies (Jarillo et al. 2017; Villaluenga, Vidal, and Cao-García 2020; Villaluenga and Cao-García 2022; Villaluenga, Brunete, and Cao-García 2023), and are derived just below.

### Extension of the Tonks-McGhee-vonHippel (TGH) model to the kinetics of ligand mode transformations

Here, we extend the binding site counting procedure to apply it to ligands that can bind in different modes (i.e., to different numbers of binding sites, as is the case for SSB) and to account for transitions between these binding modes while bound to the polymer. Within this frame, the polymer is modelled as a linear lattice with *N* identical repeated units. When a ligand molecule binds to the polymer, it covers *m* consecutive units, thereby making them inaccessible to another ligand. Thus, a free ligand binding site consists of any *m* consecutive free lattice units. One must compute the number of binding possibilities, which gives the probability of finding a binding region of the size required to accommodate a ligand molecule in a partially covered polymer. Counting is not straightforward when the ligand binds to more than one binding site because binding one ligand typically suppresses more than one binding possibility.

We define the coverage *c* of a polymer as the fraction of units covered by the ligands. Here, we consider that only one type of ligand can be bound in two modes, denoted as α and β. In mode α, the ligand binds *m*_α_ units, while in mode β, the ligand binds *m*_β_ units of the polymer (*m*_α_ < *m*_β_). The partial and total coverages are

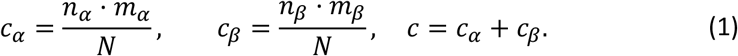

where *n*_*α*_ is the number of ligands on mode α bound to the polymer and *n*_*β*_ is the number of ligands on mode β bound to the polymer. We can define a mean effective binding mode size as

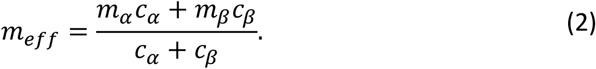

The binding of an SSB in solution to the DNA in mode *α* requires *m*_*α*_ free nucleotides, while the conversion of an SSB bound in the mode *α* to mode *β* requires *m*_*β*_ − *m*_*α*_ adjacent free nucleotides. See Fig. 1 for a graphical representation. These requirements inhibit the SSB binding (to *α* mode) and binding mode conversion (from *α* to *β* mode), leading to a decrease in the effective rates of these processes as the coverage increases. See Supplementary Information for detailed equations and their derivation.

The binding of the ligands implies a relative change of the end-to-end distance (per nucleotide) of the SSB-ssDNA complex Δ*x* (with respect to the naked ssDNA). We compute the net effect of the coverage with ligands as

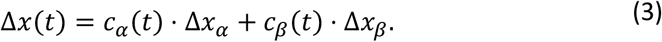

This equation links the evolution of the coverages with the evolution of the (experimentally measured) end-to-end distance through the two constant parameters Δ*x*_*α*_ and Δ*x*_*β*_.

Here, we fit our model (described below) to previous experimental results of the total coverage *c* and the mean effective mode *m*_*eff*_ on the binding to single-stranded DNA (ssDNA) of HmtSSB (Morin et al. 2017), and on the net effect on the end-to-end distance of the SSB-ssDNA complex Δ*x* for HmtSSB (Morin et al. 2017) (case with 50mM NaCl and 4 mM MgCl_2_ in the buffer) and EcoSSB (Naufer et al. 2021). The data studied here from Ref. (Morin et al. 2017) was taken at a tension of 5pN, while the data from Ref. (Naufer et al. 2021) was taken at a tension of 12pN. Rates and equilibrium constants depend on the polymer tension (Kumar and Li 2010; Jarillo et al. 2017). We chose this fixed tension for each data set because were the values with more previous experimental data available and because here we aimed to concentrate our efforts on the understanding of the kinetic effects of crowding.

#### Box 1. Kinetic equations for the two-mode binding case

The SSB binding process to ssDNA follows the reaction

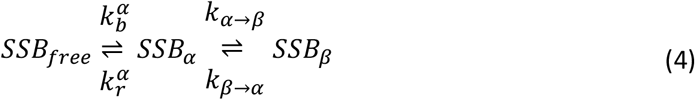

where the SSB first binds to the ssDNA at the lower binding mode *α* (covering *m*_*α*_ nucleotides), and after, it can have a transition to the higher binding mode _*β*_ (covering *m*_*β*_ nucleotides). The kinetic equations for the number of bound ligands in each mode (*n*_*α*_ and *n*_*β*_) can be expressed as

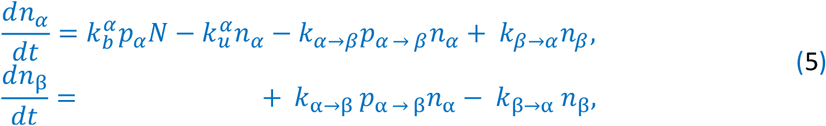

where 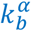 is the binding rate of the *α* mode ligand (per available binding site), 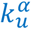 is the unboundrate of *α* mode ligand. *K*_*α*_→β gives the transition rate from the _*α*_ to the *β* mode (when possible), while *K*_*β→α*_ gives the rate of the reverse mode conversion. The binding rate 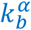 is multiplied by the number of binding locations available *p*_*α*_*N*, where *N* is the number of nucleotides in the ssDNA, and *p*_*α*_ is the probability that a nucleotide starts a free section of length *m*_*α*_, required for the binding in mode *α*. See Fig. 1A. Thus,

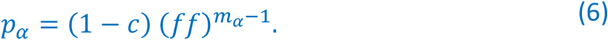

where *c* is the total coverage, given in terms of the partial coverages (*c*_*α*_ and *c*_*β*_) 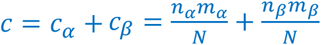. Then, 1 − *c* is the probability that a nucleotide (the first one) is free. *(ff)* is the conditional probability that, given a free nucleotide, the next-to-the-right nucleotide is also free (Villaluenga, Vidal, and Cao-García 2020). Thus, *p*_*α*_ gives the probability that a randomly chosen nucleotide is free and followed by *m*_*α*_ − 1 free nucleotides, forming thus a group of at least *m*_*α*_ free nucleotides (where a SSB can be bound in the *α* mode). The conditional probability *(ff)* is determined by the partial coverages,

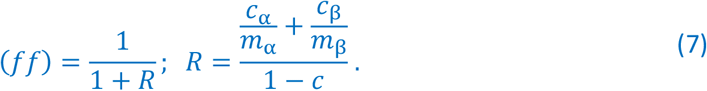

Meanwhile, the naked chain transition rate *K*_*α→β*_ is corrected in the kinetic equations, Eq. (5), by the probability

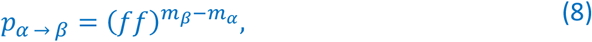

to have *m*_*β*_ − *m*_*α*_ free contiguous binding sites, which are required for the mode transformation. See Supplementary Information for details of the derivation.

The unbinding of the *α* mode does not depend on the coverage. Neither the transformation from mode β to mode α, which is always possible because the resulting ligand is shorter. Therefore, these two reversed rates do not present corrections in the kinetic equations, Eq. (5) [assuming the absence of cooperative interactions (Villaluenga and Cao-García 2022)].

The binding rate is proportional to the concentration of the ligand, [SSB], *i.e*., 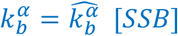. The quotients of the binding rates give the equilibrium constants for the binding in the lower mode 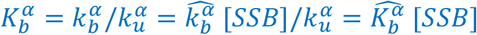 and for the mode transformation *K*_*α→β*_ = *K*_*α→β*_/*K*_*β→α*_.

## RESULTS

### Ligand crowding in DNA inhibits SSB binding

Experimental results in Ref. (Morin et al. 2017) demonstrated that HmtSSB binds to DNA with high affinity, as it remains bound even in the absence of HmtSSB in the buffer. Nevertheless, the DNA was not fully covered by HmtSSB, with observed coverages ranging from *c* = 0.80 and *c* = 0.95 (i.e., 80% to 95% coverage; see points in Fig. 2A). [The experimental coverages were obtained from force-extension curves using the mechanical models described in Ref. (Jarillo et al. 2017).] We explain this incomplete coverage by considering that each SSB binds to multiple nucleotides, and random binding creates free gaps of variable length, some too short to accommodate additional SSBs. Figure 1A provides a conceptual visualization of this crowding-inhibited binding. The detailed equations of the model are given in the Methods Section, Supplementary Information, and Refs. (McGhee and von Hippel 1974; Villaluenga, Vidal, and Cao-García 2020). Here, we assess how well the crowding-inhibited binding model explains the experimental results for HmtSSB binding to DNA reported in Ref. (Morin et al. 2017).

**Figure 2.**
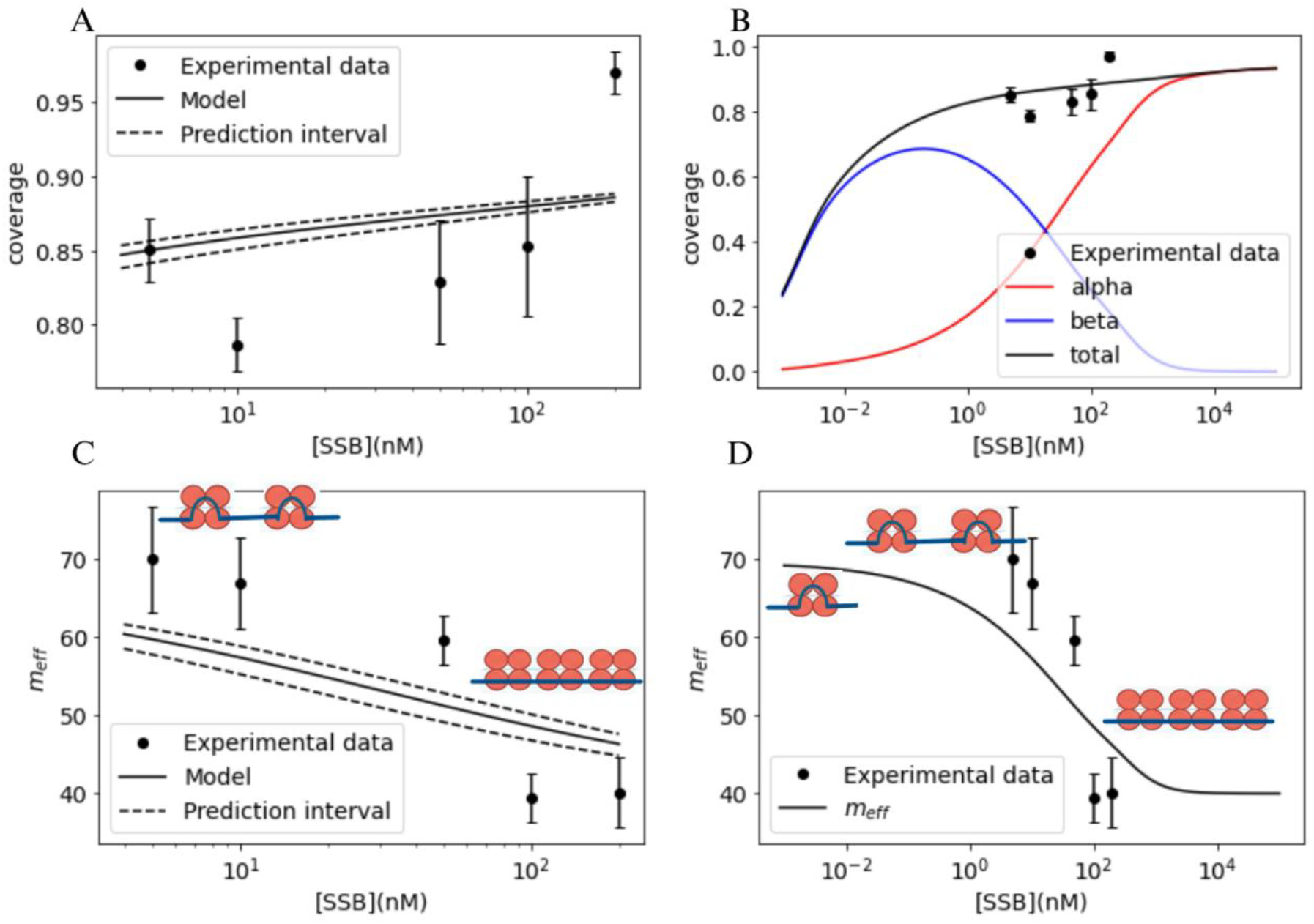
Crowding inhibition of binding and mode transformation explain equilibrium experimental data. (points) of HmtSSB binding to ssDNA reported in Ref. (Morin et al., 2017) (at low forces, 5pN, in a solution of 50mM NaCl and 4 mM MgCl_2_). Model fit is shown (black line) with a prediction confidence interval of 68 % (dashed line). Total coverage as a function of SSB concentration in solution [SSB] is shown for the experimental interval (Panel A) and extrapolated to a broader [SSB] interval (Panel B). Panel B also shows the model-predicted partial coverages due to the low (α, with m_α_ = 40) and high (β, with m_β_ = 70) modes. The effective mode size as a function of [SSB] in the experimental interval is shown in Panel C and extrapolated to a broader [SSB] interval in Panel D. Table 1 shows the values of the model parameters obtained from the fit shown here. Panels show how increasing the [SSB] concentration in the external solution increases coverage (without attaining complete coverage) and shifts the prevailing binding mode from the higher binding mode (β) to the lower binding mode (α).

The equilibrium model considers the two binding modes (*m*_*α*_ = 40 and *m*_*β*_ = 70) and depends on two parameters: the equilibrium constant of the lower mode binding 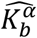 (or equivalently, the value at a reference SSB concentration, 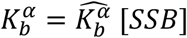 and the equilibrium constant of the mode transformation *K*_*α*_→_*β*_. These two parameters are determined simultaneously fittting the coverage (Fig. 2A) and the effective mean mode size *m*_*eff*_ (Fig. 2C), see Table 1. (For details, see Supplementary Information). Fig. 2B and 2D also show the model’s extrapolation to lower and higher SSB concentrations.

**Table 1:**
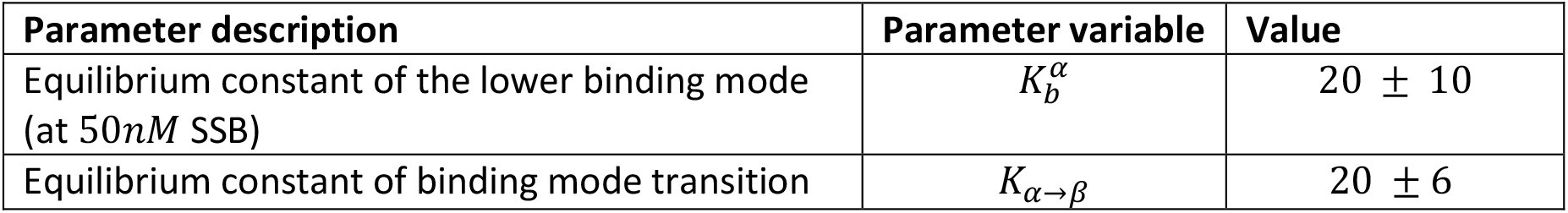
Parameters obtained from the fit of the equilibrium model to the experimental data of HmtSSB binding to DNA in Ref. (Morin et al. 2017) at low forces, 5pN, for a solution with 50mM NaCl and 4 mM MgCl_2_. (m_α_ = 40 and m_β_ = 70.) See fit in Fig. 2.

Therefore, our crowding inhibition model can conceal high binding affinities, 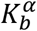 from 20 (at 50*nM* Table 1) to 80 (at 200*nM*) with partial coverages, 80 to 90% (see Fig. 2A). In contrast, a simpler Langmuir-like model would predict for this range of affinities coverages in the range of 95 to 99%.

### Crowding in DNA inhibits the transition to higher SSB binding modes

Experimental data of Ref. (Morin et al. 2017) for HmtSSB shows that the mean effective mode *m*_*eff*_ depends on the SSB concentration in solution, see points in Fig. 2C. This effect can be explained by the inhibition of the transition to higher modes induced by crowding, as shown in Fig. 1B. In a low-covered DNA molecule, there are many available free binding sites to accommodate ligand molecules, and the conversion between the low mode α and the high mode β occurs easily. Figure 1B (left) illustrates this behavior. In our model, the increase in DNA coverage significantly reduces the number of accessible binding locations of appropriate size, inhibiting the conversion from low site-size mode α to high site-size mode β. Figure 1B (right) illustrates this behavior. Our model predicts that the high-binding mode β (70 nt) dominates at low SSB concentrations, while the low-binding mode α (40 nt) prevails at high SSB concentrations (see Figure 2B), in agreement with observations. This decrease in the mean effective binding mode size increases the maximum attainable coverage, as discussed below.

As explained above, the joint fit of the coverage and the mean effective mode as a function of [SSB] provides the equilibrium constants for the lower mode binding 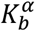 (or 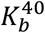) and for the binding mode transformation *K*_*α→β*_ (or *K*_40→70_). Fig. 2 shows the fit, and Table 1 shows the resulting equilibrium constant values.

### Ligand crowding inhibition explains experimentally observed SSB concentration dependence of the coverage

Previous experimental results have shown that DNA coverage by HmtSSB (Morin et al. 2017) increases from 80 to 95 % when HmtSSB concentration in solution increases from 5 nM to 200 nM. See Fig. 2. Our model explains this rise in coverage with SSB concentration with a reasonable agreement (see Figures 2A and 2B). In our model, the observed increase in coverage arises from the effective decrease in mode size *m*_*eff*_. As we discussed in the previous point, the increase in [SSB] leads to a shift from the high binding mode (*m*β = 70) to the low binding mode (*m*α = 40), Fig. 2C and 2D. Ligands in the lower binding mode can bind to smaller gaps, leading to higher coverages (Villaluenga, Vidal, and Cao-García 2020; Villaluenga, Brunete, and Cao-García 2023). Therefore, the shift towards the low binding mode with increasing [SSB] shown in Fig. 2C implies the increase in the coverage with increasing [SSB] shown in Fig. 2A.

### Crowding-induced changes in effective rate constants explain bimodal kinetics

Experimental data of HmtSSB binding to ssDNA showed bimodal kinetics in the SSB concentrations close to the transition between low and high binding mode in equilibrium (the region with experimental data in Fig. 2)(Morin et al. 2017). This bimodal binding kinetics consists of a rapid shortening of the end-to-end distance of the SSB-ssDNA complex, followed by a slow expansion. See Fig. 3. Our model explains the initial shrinking kinetics as a process dominated by the binding in the lower (alpha) mode and its fast conversion to the higher (beta) mode (left part of Panels B and C of Fig. 3). Later, binding and conversion are inhibited by the increasing coverage, leading to a slowly expanding kinetics dominated by the slow conversion of ligand from the higher binding mode to the lower binding mode, together with a slow increase in coverage (see right part of Panels in Fig. 3). This increase in the crowding-induced inhibition of the low to high mode transition shifts the equilibrium to the lower mode, see Fig. 3D. (Fig. 1B shows the mechanism of crowding-induced inhibition of transitions from low to high binding mode.) Fig. 3D compares the ratio of the effective rates and the ratio of the partial coverages. They match in equilibrium. However, as the effective rates depend on the partial coverages, this introduces feedback. The feedback makes the final ratios to reach vary as the coverage ratio goes to it, leading to a slow approach to a shifting equilibrium.

### Crowding also accounts for EcoSSB binding kinetics (and the changes reported for the effective rates)

Experimental data on the binding, unbinding, and rebinding kinetics of EcoSSB on ssDNA (Naufer et al. 2021) reveal an interesting behavior in the extension kinetics. See Fig. 4A. The addition of EcoSSB to the buffer induces a shortening of the end-to-end distance of ssDNA due to the formation of an EcoSSB-ssDNA complex (see Fig. 4A left). Surprisingly, removing EcoSSB results in a further contraction (Fig. 4A middle), which is reverted when EcoSSB is returned to the solution (Fig. 4A right). This surprising further contraction (induced by the EcoSSB remotion from the buffer) can be understood as an effect of allowing the transition from lower to higher binding modes, facilitated by the unbinding of a fraction of the ligands bound in the lower mode. This unbinding reduces the crowding-induced inhibition of the transition to higher modes (see Fig. 1B for a representation of the process).

**Figure 3.**
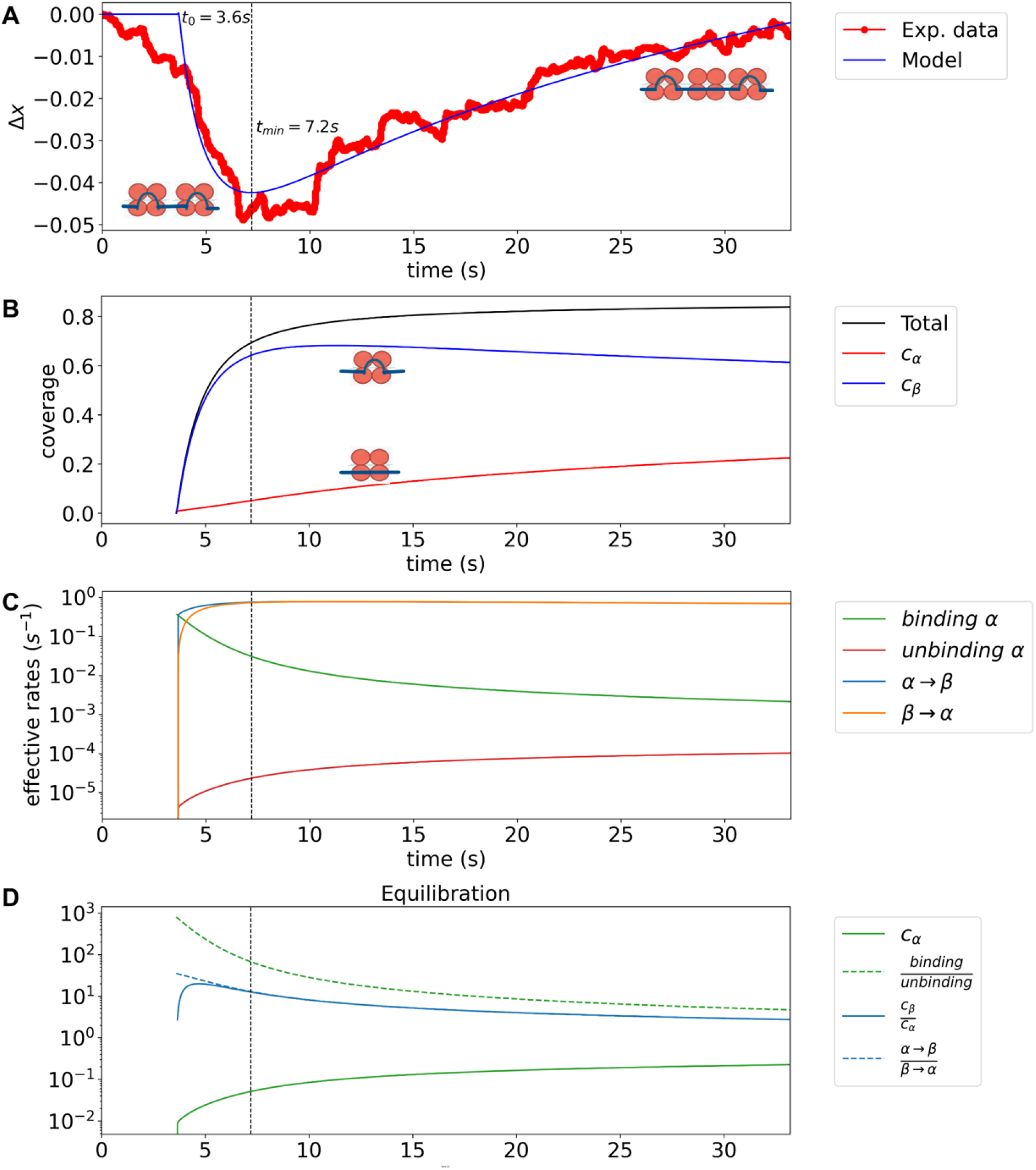
The proposed crowding inhibition model reproduces HmtSSB bimodal binding kinetics. HmtSSB binding to ssDNA presents a bimodal binding kinetics with a transient shortening in the end-to-end length of the SSB-ssDNA complex (Panel A). Evolution is fitted to the two-mode kinetic model (thin blue line) using the equilibrium constants from Table 1. The results of the least-squares fit are shown in Table 2. (The effective initial binding time t_0_ = 3.6s is also fitted) Panel B shows the evolution of the model’s partial and total coverage. Panel B implies that the bimodal kinetics can be explained by a fast binding in the lower α mode and conversion to the higher β mode, followed by a slow conversion from β to α mode as coverage increases with time. Panel C compares the effective rates of the alpha coverage kinetic equation. Effective rates include the crowding inhibition corrections. Panel C shows that α binding dominates over release. In contrast, mode conversion from α to β fastly reaches equilibrium. These results are even clearer in Panel D. Panel D compares the ratio of the coverage-dependent effective rates with the ratio of the partial coverages. They match in equilibrium. Panel D shows the rapid equilibration between modes and the slow increase in the alpha coverage towards its equilibrium value. The lower binding mode, α, binds to mα = 40 nucleotides, while the higher binding mode, β, binds to m_β_ = 70 nucleotides. Experimental values are detailed in the caption of Table 2.

**Figure 4.**
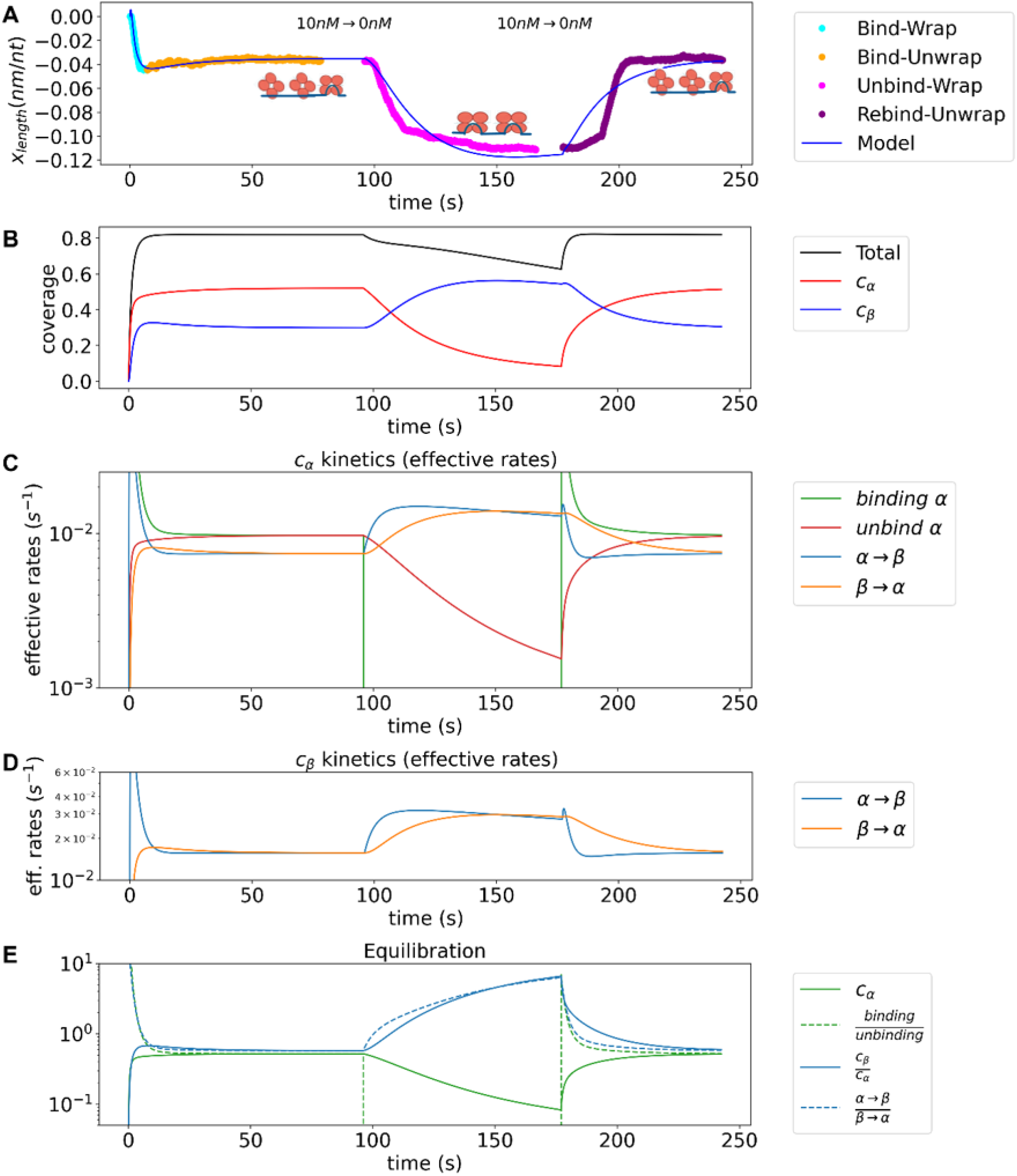
The proposed crowding inhibition model with two modes describes the binding, unbinding, rebinding kinetics of EcoSSB on ssDNA. Panel A shows the experimental (Naufer et al. 2021) extension of the SSB-ssDNA complex as a function of time (while EcoSSB is added, removed, and added again to the solution) compared with our model with two binding modes for EcoSSB (using the best-fit values given in Table 3). Panel B shows the evolution of the partial coverages in the α mode (low, m_α_ = 8) and β mode (medium, m_α_ = 17), and the total coverage provided by the model. Panel B also shows that the withdrawal of EcoSSB from the buffer results in the release of some of the EcoSSB bound in the α mode, reducing coverage and thereby allowing the conversion of other EcoSSB bound in the α mode to the β mode. Panel C shows the evolution of the effective rates of the different processes involved in the kinetic equation of the alpha mode. Further clarifying the dominant process in each time interval. Panel D shows the evolution of the effective rates for the different processes involved in the β-mode kinetic equation. Panel E compares the ratio of the effective rates to the ratio of the partial coverages. Experimental data are from Fig. 3A of Ref. (Naufer et al. 2021), which was taken at a tension of 12 pN.

Fig. 4 shows the comparison of the experimental data and the (two mode) crowded-inhibition model for end-to-end extension (Panel A), the model evolution of the partial and total coverage (Panel B), the comparison of the terms in the evolution equations (expressed in coverage decrease per unit of time, Panel C), and the evolution of the coverage rates and its value in equilibrium computed with the rate of effective rates (Panel D). The coverage-dependence of the effective rates also gives rise to an evolution of the effective rates ratio, leading to an evolving equilibrium point to reach, as shown in the last panel of Fig. 4.

**Table 2:** Parameter values of the two binding mode model used in Fig. 3 to compare with experimental data (Morin et al. 2017) of HmtSSB binding to ssDNA. The binding rate of the lower mode 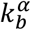 and transition rate from the higher to the lower mode K_β→α_ are the two free parameters of the model, the other two rates are fixed by the equilibrium constants (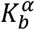and K_α→β_) taken from the previous fits to equilibrium data (Table 1 and Fig. 2) (m_α_ = 40 and m_β_ = 70). Δx is the end-to-end distance of the SSB-ssDNA complex (per nucleotide with respect to the ssDNA naked polymer). Experimental data for [SSB] = 50 nM and F = 5 pN tension, in a solution of 50mM NaCl and 4 mM MgCl_2_ from Ref. (Morin et al. 2017).

| Parameter description | Parameter variable | Value |
| --- | --- | --- |
| Binding rate of lower mode | $k_b^\alpha$ | $9.2 \cdot 10^{-3} \text{ s}^{-1}$ |
| Unbinding rate of lower mode | $k_u^\alpha$ | $9.2 \cdot 10^{-3} \text{ s}^{-1} / K_b^\alpha$<br>$= 4.6 \cdot 10^{-4} \text{ s}^{-1}$ |
| Transition rate from lower to higher mode | $k_{\alpha \rightarrow \beta}$ | $(2.0 \text{ s}^{-1}) \cdot K_{\alpha \rightarrow \beta}$<br>$= 40 \text{ s}^{-1}$ |
| Transition rate from higher to lower mode | $k_{\beta \rightarrow \alpha}$ | $2.0 \text{ s}^{-1}$ |
| α-mode induced relative change in $\Delta x$ | $\Delta x_\alpha$ | $0.22 \text{ nm/nt}$ |
| β-mode induced relative change in $\Delta x$ | $\Delta x_\beta$ | $-0.08 \text{ nm/nt}$ |

**Table 3:** Parameter values of the two binding modes model obtained from the least square fit to the experimental data of EcoSSB binding to ssDNA from Ref. (Naufer et al. 2021) at 12 pN and binding at [EcoSSB] = 10nM. Fit shown in Fig. 4A. (The same values were also obtained with Bayesian inference, see Supplementary Information.) The modes α and β, are assumed [as in Ref. (Naufer et al. 2021)] to bind to 8 and 17 nucleotides, respectively (i.e., m_α_ = 8 and m_β_ = 17). All other parameters are free for this case. Δx is the end-to-end distance of the SSB-ssDNA complex (per nucleotide with respect to the ssDNA naked polymer). Results are compatible with previous experimental results in Fig. 3A of Ref. (Suksombat et al. 2015), which reported values of Δx_17_ ~ −0.2 nm/nt for the 17-mode of E. Coli SSB at 12 pN.

| Parameter description | Parameter variable | Values |
| --- | --- | --- |
| Binding rate of α-mode | $k_b^\alpha = k_b^8$ | $9.1 \times 10^{-2} \text{ s}^{-1}$ |
| Unbinding rate of α-mode | $k_u^\alpha = k_u^8$ | $1.9 \times 10^{-2} \text{ s}^{-1}$ |
| Transition rate from $\alpha$ to β-mode | $k_{\alpha \rightarrow \beta} = k_{8 \rightarrow 17}$ | $0.41 \text{ s}^{-1}$ |
| Transition rate from $\beta$ to α-mode | $k_{\beta \rightarrow \alpha} = k_{17 \rightarrow 8}$ | $5.3 \times 10^{-2} \text{ s}^{-1}$ |
| α-mode induced change in $\Delta x$ | $\Delta x_\alpha = \Delta x_8$ | $5.9 \times 10^{-2} \text{ nm/nt}$ |
| β-mode induced change in $\Delta x$ | $\Delta x_\beta = \Delta x_{17}$ | $-0.22 \text{ nm/nt}$ |

EcoSSB is known to bind ssDNA in several modes, with binding to 8, 17, 35, and higher numbers of nucleotides (Suksombat et al. 2015; Naufer et al. 2021). Here, we show that our model for crowded binding with two modes, EcoSSB_8_ and EcoSSB_17_, can explain the main features observed in the experimental kinetic data for EcoSSB binding to ssDNA on Ref. (Naufer et al. 2021). Also, in Ref. (Naufer et al. 2021), the same EcoSSB modes were considered, as they dominate under buffer and tension conditions and are supported by compatible observed end-to-end distance shrinkage (Suksombat et al. 2015). See Table 3 and Fig. 4, for the parameter values and plot resulting from the fit.

The binding kinetics show a bimodal kinetics with a fast shortening of the end-to-end distance of the EcoSSB-ssDNA complex, followed by a slow expansion (left part of Panel A of Fig. 4). (Analogous to the one described above for HmtSSB.) We show that our model can explain the first part of this bimodal kinetics (thick blue line in Fig. 4A) as a fast binding in the EcoSSB_8_ mode, simultaneous with a fast conversion to the EcoSSB_17_ mode (wrapping). This fast first part leads to high coverage, increasing the crowding-induced inhibition of binding and wrapping (EcoSSB_8_ to EcoSSB_17_ conversion). This crowding-induced inhibition initiates a second part of the kinetics, a slow phase combining slow binding in the lower mode, EcoSSB8, and a slow conversion from the higher to the lower mode, EcoSSB_17_ to EcoSSB_8_. As the coverage slowly increases, it also increases the inhibition of the transition to the higher mode, shifting the equilibrium towards the lower mode. (See the left part of all panels in Fig. 4.)

The experimental unbinding kinetics (induced by removing the SSB from the solution, *[SSB]* = 0 *nM*) shows an intriguing shrinkage (middle part of Panel A of Fig. 4) (Naufer et al. 2021). The crowding model we propose here can explain this shrinking as the net unbinding of some ligands bound in the low mode (EcoSSB_8_), which allows the conversion of other EcoSSB_8_ to the higher EcoSSB_17_. (See the middle part of all panels in Fig. 4.) This process leads to shrinking provided the conversion is faster than the unbinding, and the higher mode exhibits greater shrinking. See Table 3.

The rebinding kinetics has an experimental return to the previous bound equilibrium (right part of Panel A of Fig. 4), which our model can describe as an increased crowding that inhibits the transitions to the higher mode (EcoSSB_17_), leading to a decay to the lower mode (EcoSSB_8_) until the previous EcoSSB_8_ and EcoSSB_17_ binding equilibrium is restored. See Fig. 4.

These implications of the crowding-induced inhibition model are compatible with the phenomenological explanation already provided in Ref. (Naufer et al. 2021). Our model adds explicit expressions that account for the crowding inhibition of SSB binding and transition to higher modes, as effective rates, during the complete binding, unbinding, and rebinding experiment. Thus, our crowding inhibition model provides the required changes in the effective rates to reproduce the experimentally observed kinetics.

## DISCUSSION AND CONCLUSIONS

The crowded-inhibited binding model presented here accurately describes the key features of the SSB binding equilibrium and kinetics observed experimentally, for both HmtSSB and EcoSSB (Morin et al. 2017; Naufer et al. 2021). In this model, crowding inhibition arises from the scarcity of consecutive free binding sites required for binding or for transitions to higher binding modes, as quantified using the Tonks-McGhee-von Hippel (TGH) counting procedure. The model reproduces the experimentally observed crowding inhibition of SSB ligand binding and SSB ligand conversion to higher binding modes. The model provides explicit expressions for the coverage-dependent effective binding and transformation rates, which fit the experimentally observed kinetics.

The model can also be applied to other multimode ligands binding to polymers, allowing prediction of the effects of crowding on ligand binding and mode selection. While cooperativity has not been included here—since the binding modes considered do not exhibit significant cooperativity—it can be incorporated by combining the present kinetic equations with those from Ref. (Villaluenga and Cao-García 2022). Extending the model to include cooperativity will allow to explore the accuracy of the resulting framework to explain also the similar multimode kinetics observed for L1-ORF1p nucleoprotein and for T4 gene 32 protein in Refs. (Cashen et al. 2024, 2023).

Similarly, competition with other ligands can be modeled using the approaches described in Ref. (Villaluenga, Brunete, and Cao-García 2023). Incorporating interactions between non-nearest-neighbors (Bonde et al. 2024) would require further development or resorting to more computationally intensive molecular dynamics simulations. However, in optical or magnetic tweezers experiments, where ssDNA is under tension, such non-nearest-neighbor interactions are energetically disfavored due to the associated large shortening of the SSB-ssDNA complex.

Another aspect to consider is the potential impact of SSB condensate formation (Gábor M. Harami et al. 2020; Gabor M. Harami and Neuman 2022; Kozlov et al. 2022; Bonde et al. 2024). SSB condensates could, in principle, form on ssDNA and/or in solution. On SSB, while SSBs bound to ssDNA could be compacted via non-nearest-neighbor interactions, such interactions are energetically unfavorable under the tension applied in optical and magnetic tweezers experiments (as commented above). In solution, SSB condensates are reported to occur and are inhibited by ssDNA (Bonde et al. 2024). The formation of SSB condensates in solution would reduce the availability of free SSBs to bind ssDNA, requiring a correction to the effective free SSB concentration in the binding kinetics.

## ACKNOWLEDGEMENTS

We acknowledge Mark C. Williams and his team for providing the experimental data on the kinetics of EcoSSB binding to DNA from their work (Naufer et al. 2021). We thank the members of our laboratories for useful discussions during this work and our previous work on HmtSSB (Morin et al. 2017).

This study was supported by MCIN/AEI/10.13039/501100011033 (grants PGC2018-099341-B-I00 and PID2021-126755NB-I00 to B.I., RTI2018-095802-B-I00 and PID2023-148319NB-I00 to F.J.C.G.); and Universidad Complutense de Madrid (UCM) (PR12/24-31558 to F.J.C.G).

English and sentence structure improved with Grammarly. Structure of sentences and paragraphs improved with Nature Research Assistant. The authors carefully checked that the improved sentences and paragraphs kept the original intended meaning. From the text abstract, Gemini provided a first graphical abstract manually corrected by the authors on Powerpoint.

## Authors contributions

Alberto Perez-Mugia (Formal analysis [supporting], Software [supporting], Visualization [supporting], Writing – original draft [supporting])

Bruno Marcos (Formal analysis [supporting], Resources [supporting], Validation [supporting], Software [supporting], Visualization [supporting], Writing – original draft [supporting])

Juan P. García-Villaluenga (Methodology [supporting], Writing – original draft [supporting])

Borja Ibarra (Writing – review & editing [supporting])

Francisco J. Cao-García (Conceptualization [lead], Methodology [lead], Formal analysis [lead], Software [lead], Resources [lead], Validation [lead], Visualization [lead], Funding acquisition [lead], Writing – original draft/review & editing [lead])

## Data availability statement

The data underlying this article will be shared on reasonable request to the corresponding author.

## Notes

### Competing Interest Statement

The authors have declared no competing interest.

