## Supplementary Information for "Crowding on DNA modulates SSB protein binding mode kinetics"

#### 1. Transitions between two binding modes to a long polymer (lattice model)

Here, we extend the binding site counting procedure to apply it to ligands that can bind in different modes (i.e., to different numbers of binding sites) and transition between these modes while bound to the polymer.

From a physical perspective, a bound ligand molecule occupies a certain number of consecutive units. Some ligands can bind a polymer in multiple modes, differing in the number of occluded polymer units. Coverage $c$ is defined as the fraction of units covered by the ligands. Ligands in mode $i$ cover a fraction of the polymer:

$$c_i = \frac{n_i \cdot m_i}{N}, \tag{S1}$$

where $n_i$ is the number of ligands on mode $i$ bound to the polymer, and $m_i$ is the number of polymer units covered by the ligand on mode $i$, and $N$ is the total number of repeating units in the polymer. Note that each unit can be in only two states: bound to the ligand or free. Then, the total fraction of the polymer that is bound to ligands is given by,

$$c = \sum_{i=1}^{k} c_i, \tag{S2}$$

where $k$ is the number of different modes of ligands (Villaluenga, Brunete, and Cao-García 2023).

In our approach, inspired by the McGhee and von Hippel procedure (McGhee and von Hippel 1974, 1976), the main object to compute is the number of binding possibilities, given by the probability of finding a binding region of the required size to accommodate a ligand molecule in a partially covered polymer.

Here, we assume that there is only one type of ligand that can bind in two modes, denoted α and β. In mode α, the ligand binds $m_\alpha$ units, while mode β the ligand binds $m_\beta$ units of the polymer. Therefore, the partial and total coverages are given by

$$c_\alpha = \frac{n_\alpha \cdot m_\alpha}{N}, \qquad c_\beta = \frac{n_\beta \cdot m_\beta}{N}, \quad c = c_\alpha + c_\beta. \tag{S3}$$

A useful relationship is obtained by restating the probability of finding an empty site, $1-c$, in terms of the state of the previous site and the conditional probabilities for the current site given the state of the previous one. The previous site could be empty (with probability $1-c$), occupied by the last element of a ligand of mode $\alpha$ (with probability $c_\alpha/m_\alpha$) or occupied by the last element of a ligand of mode $\beta$ (with probability $c_\beta/m_\beta$). Note that other states of the previous site are not possible as they would not allow the current site to be empty. These considerations allow us to write the following consistency equation (McGhee and von Hippel 1974, 1976; Villaluenga, Brunete, and Cao-García 2023)

$$(1-c)(ff)+\frac{c_{\alpha}}{m_{\alpha}}\left(b_{m_{\alpha}}^{\alpha}f\right)+\frac{c_{\beta}}{m_{\beta}}\left(b_{m_{\beta}}^{\beta}f\right)=(1-c), \tag{S4}$$

where $(ff)$ is the probability, given a free unit, that another free unit lies to the immediate right; $\left(b_{m_i}^{i}f\right)$ is the probability, given the right end of a bound ligand of mode $i$, that a free unit lies to the immediate right.

For noncooperative ligands, there are neither attractive nor repulsive interactions between bound ligands. This absence of inter-ligand interactions implies that the probability of having an empty site is independent of the state of the previous site, i.e., $\left(b_{m_i}^{i}f\right)=(ff)$, and Eq. (S4) becomes

$$\frac{1-(ff)}{(ff)}=R\,, \tag{S5}$$

with $R$ defined as

$$R=\frac{\left(\frac{c_{\mathrm{\alpha}}}{m_{\mathrm{\alpha}}}+\frac{c_{\mathrm{\beta}}}{m_{\mathrm{\beta}}}\right)}{1-c}. \tag{S6}$$

One can solve these equations to obtain $(ff)$ as a function of the partial coverages,

$$(ff)=\frac{1}{1+R}=\frac{1-c_{\mathrm{\alpha}}-c_{\mathrm{\beta}}}{1-c_{\mathrm{\alpha}}-c_{\mathrm{\beta}}+\frac{c_{\mathrm{\alpha}}}{m_{\mathrm{\alpha}}}+\frac{c_{\mathrm{\beta}}}{m_{\mathrm{\beta}}}}, \tag{S7}$$

We also consider that these two modes can transform into each other while they are bound to the polymer. If we assume, without loss of generality, that $m_{\mathrm{\alpha}}<m_{\mathrm{\beta}}$, the transformation from mode α to mode β is only possible if there are $m_{\mathrm{\beta}}-m_{\mathrm{\alpha}}$ free units next to an α ligand. Accordingly, the probability of having these free units is given by,

$$p_{\mathrm{\alpha}\to\mathrm{\beta}}=\left(b_{m_{\mathrm{\alpha}}}^{\mathrm{\alpha}}f\right)(ff)^{m_{\mathrm{\beta}}-m_{\mathrm{\alpha}}-1}. \tag{S8}$$

The transformation from mode β to mode α is always possible because the resulting ligand is shorter, i.e., $p_{\mathrm{\beta}\to\mathrm{\alpha}}=1$.

#### 2. Kinetic equations for a ligand with transitions between two binding modes to a long polymer

SSBs may bind ssDNA in multiple modes that differ in the number of nucleotides of ssDNA bound. Experimental data show that Human mitochondrial SSB (HmtSSB) binds ssDNA in at least two modes $(SSB)_{\mathrm{\alpha}}$ and $(SSB)_{\mathrm{\beta}}$, which bind $m_{\alpha}\simeq 40$ and $m_{\beta}\simeq 70$ nucleotides of ssDNA (Morin et al. 2017). Escherichia coli SSB (EcoSSB) may bind ssDNA in several modes, which we denote as (SSB)$_8$, (SSB)$_{17}$, (SSB)$_{35}$, (SSB)$_{56}$, and (SSB)$_{65}$, where the subscripts here indicate the average number of ssDNA nucleotides occluded per SSB protein (Kuznetsov et al. 2006; Antony et al. 2013; Suksombat et al. 2015; Naufer et al. 2021). At higher tension, as in the data we analyze here ($12pN$), only the two lower modes of EcoSSB are expected to be relevant (Naufer et al. 2021). On both SSB-ssDNA systems, the prevalent modes depend on the SSB concentration in the solution.

We consider that a ligand molecule can bind to the polymer in two distinct modes, differing in the number of occluded units (Morin et al. 2017; Naufer et al. 2021). Therefore, counting also the unbound state, a ligand can be in three states: a free ligand in solution (free state), a ligand bound to the polymer occluding $m_\alpha$ units (state or mode α), and a ligand bound to the polymer occluding $m_\beta$ (state or mode β). The process of ligand binding starts with the incorporation of a ligand from the free state to the state α (with a binding rate $k_b^\alpha$), which is the state with smaller occluded units ($m_\alpha < m_\beta$). (The transformation is reversible with unbinding rates $k_u^\alpha$.) Following the binding in state α, the ligand can be transformed from state α to a state β with a transformation rate $k_{\alpha\to\beta}$. The ligand is also allowed to evolve from state β to state α with transformation rate $k_{\beta\to\alpha}$.

$$SSB_{free} \underset{k_u^\alpha}{\overset{k_b^\alpha}{\rightleftharpoons}} SSB_\alpha \underset{k_{\beta\to\alpha}}{\overset{k_{\alpha\to\beta}}{\rightleftharpoons}} SSB_\beta \tag{S9}$$

The kinetic equations with transitions between binding modes are obtained here by generalizing the previous results for the binding of two ligand modes to a linear polymer (Villaluenga, Brunete, and Cao-García 2023). Assuming that $m_\alpha < m_\beta$, the transition $\alpha$ to $\beta$ requires to have $m_\beta - m_\alpha$ free nearby binding sites, which happens with probability $p_{\alpha\to\beta}$, which enters as a factor correcting the naked chain transition rate $k_{\alpha\to\beta}$. Thus, the kinetic equations describing the time variation in the number of ligands bounded to the polymer is given by

$$\begin{aligned}\frac{dn_\alpha}{dt} &= k_b^\alpha\, p_\alpha N - k_u^\alpha\, n_\alpha - k_{\alpha\to\beta}\, p_{\alpha\to\beta}\, n_\alpha + k_{\beta\to\alpha}\, n_\beta,\\ \frac{dn_\beta}{dt} &= \qquad\qquad\qquad\qquad + k_{\alpha\to\beta}\, p_{\alpha\to\beta}\, n_\alpha - k_{\beta\to\alpha}\, n_\beta.\end{aligned} \tag{S10}$$

$p_\alpha$ is the probability that a binding site of length $m_\alpha$ starts at a position chosen at random. Thus, $p_\alpha N$ is the number of binding sites for a ligand in mode $\alpha$ (for a long polymer where $N - m_\alpha \simeq N$), expressed in previous publications as $p_\alpha N = (n+1)\bar{s}_\alpha$, where $\bar{s}_\alpha$ is the average number of free binding sites per gap in the polymer (Villaluenga, Vidal, and Cao-García 2020; Villaluenga and Cao-García 2022). The number of bound ligands is $n$, and the number of gaps is $n + 1 \simeq n$ for large values (McGhee and von Hippel 1974, 1976). The derivation of the average number of free binding sites per gap $\bar{s}_\alpha$ can be found in Ref. (Villaluenga, Brunete, and Cao-García 2023), which gives, for the noncooperative case,

$$\begin{aligned}\bar{s}_\alpha &= \left[\sum_{\lambda=1}^{k} \frac{c_\lambda}{c}\left(b_{m_\lambda}^\lambda f\right)\right] \frac{(ff)^{m_\alpha-1}}{\left(1-(ff)\right)^2} \left[\sum_{\nu=1}^{k} (f b_1^\nu)\right] = \\ &= \left[\sum_{\lambda=1}^{k} \frac{c_\lambda}{c}\right](ff)\frac{(ff)^{m_\alpha-1}}{\left(1-(ff)\right)^2}\left(1-(ff)\right) = \frac{(ff)^{m_\alpha}}{1-(ff)}.\end{aligned} \tag{S11}$$

In the derivation of this expression, we use that $c = \sum_{\lambda=1}^{k} c_\lambda$, and that in the noncooperative case $\left(b_{m_\lambda}^\lambda f\right) = (ff)$.

Here, as we have two binding modes, $\alpha$ and $\beta$, the total number of ligands attached to the polymer is given by $n = n_\alpha + n_\beta$. The probability is given by

$$p_\alpha = \bar{s}_\alpha \frac{n}{N} = \bar{s}_\alpha \frac{n_\alpha + n_\beta}{N} = \frac{(ff)^{m_\alpha}}{1-(ff)}\left(\frac{c_\alpha}{m_\alpha} + \frac{c_\beta}{m_\beta}\right) = \frac{(ff)^{m_\alpha}}{1-(ff)} R(1-c) = \\ = (ff)^{m_\alpha - 1}(1-c) \tag{S12}$$

where we have used the expressions for the conditional probability $(ff)$ in the noncooperative case given by Eqs. (S5), (S6), and (S7).

Following an analogous procedure, we have in the noncooperative case that the probability of the conversion from mode α to mode β is allowed, $p_{\alpha\to\beta}$ , can be simplified as

$$p_{\alpha\to\beta} = \left(b^{\alpha}_{m_\alpha} f\right)(ff)^{m_\beta - m_\alpha - 1} = (ff)^{m_\beta - m_\alpha}. \tag{S13}$$

This expression gives the probability that the transition is allowed, computing the probability that there is a nearby empty gap or at least $m_\beta - m_\alpha$ free binding sites (i.e., free nucleotides).

The kinetic equations for the number of ligands bound to the polymer, Eq. (S10), using Eqs. (S12) and (S13), is expressed as

$$\begin{aligned} \frac{dn_\alpha}{dt} &= k_b^\alpha (ff)^{m_\alpha - 1}(1-c)N - k_u^\alpha n_\alpha - k_{\alpha\to\beta}(ff)^{m_\beta - m_\alpha} n_\alpha + k_{\beta\to\alpha} n_\beta, \\ \frac{dn_\beta}{dt} &= +k_{\alpha\to\beta}(ff)^{m_\beta - m_\alpha} n_\alpha - k_{\beta\to\alpha} n_\beta. \end{aligned} \tag{S14}$$

These kinetic equations can be expressed in terms of the coverages, using Eq. (S1), as

$$\begin{aligned} \frac{dc_\alpha}{dt} &= k_b^\alpha (ff)^{m_\alpha - 1} m_\alpha (1-c) - k_u^\alpha c_\alpha - k_{\alpha\to\beta}(ff)^{m_\beta - m_\alpha} c_\alpha + k_{\beta\to\alpha} \frac{m_\alpha}{m_\beta} c_\beta, \\ \frac{dc_\beta}{dt} &= +k_{\alpha\to\beta}(ff)^{m_\beta - m_\alpha} \frac{m_\beta}{m_\alpha} c_\alpha - k_{\beta\to\alpha} c_\beta, \end{aligned} \tag{S15}$$

These kinetic equations, Eq. (S14) or (S15), allow us to model the evolution of the system.

The equilibrium constants for binding and transformation are given by

$$K_b^\alpha = \frac{k_b^\alpha}{k_u^\alpha}, \quad K_{\alpha\to\beta} = \frac{k_{\alpha\to\beta}}{k_{\beta\to\alpha}}, \tag{S16}$$

where $K_\alpha$ is the equilibrium constant that relates the binding and unbinding rate constants of the α ligand, and $K_{\alpha\to\beta}$ is the equilibrium constant of the mode transformation α to β.

The binding rate is proportional to the concentration of SSB in solution, $[SSB]$. Thus, we can express the binding rate and the binding constant as

$$\begin{aligned} k_b^\alpha &= \widehat{k_b^\alpha} \cdot [SSB], \\ K_b^\alpha &= \frac{k_b^\alpha}{k_u^\alpha} = \frac{\widehat{k_b^\alpha} \cdot [SSB]}{k_u^\alpha} = \widehat{K_b^\alpha} \cdot [SSB], \end{aligned} \tag{S17}$$

where the hat quantities, $\widehat{k_b^\alpha}$ and $\widehat{K_b^\alpha}$, express their value per unit of SSB concentration.

Additionally, for interpreting the experimental data in Ref. (Morin et al. 2017), it will be useful to define an effective mode size function as

$$m_{eff} = \frac{c_\alpha m_\alpha + c_\beta m_\beta}{c_\alpha + c_\beta}, \tag{S18}$$

plotted as a function of the concentration of SSB in solution in Fig. 2C and D of the main text.

#### 2. Equilibrium coverage for a ligand with transitions between two binding modes to a long polymer

In equilibrium, the derivatives of both partial coverages, $c_\alpha$ and $c_\beta$, vanish. Thus, the kinetic equations, Eq. (S15), give the following system of equilibrium conditions,

$$0 = k_b^\alpha (ff)^{m_\alpha - 1} m_\alpha (1-c) - k_u^\alpha c_\alpha - k_{\alpha\to\beta}(ff)^{m_\beta - m_\alpha} c_\alpha + k_{\beta\to\alpha}\frac{m_\alpha}{m_\beta} c_\beta, \tag{S19}$$

$$0 = k_{\alpha\to\beta}(ff)^{m_\beta - m_\alpha}\frac{m_\beta}{m_\alpha} c_\alpha - k_{\beta\to\alpha} c_\beta. \tag{S20}$$

The last equation can be written in a more convenient form, as an expression giving the coverage ratio

$$A \equiv \frac{c_\beta}{c_\alpha} = \frac{k_{\alpha\to\beta}}{k_{\beta\to\alpha}}(ff)^{m_\beta - m_\alpha}\frac{m_\beta}{m_\alpha} = K_{\alpha\to\beta}(ff)^{m_\beta - m_\alpha}\frac{m_\beta}{m_\alpha}. \tag{S21}$$

The conditional probability $(ff)$ is given by Eq. (S7), and in terms of the coverage ratio $A$ and the coverage with mode α, $c_\alpha$, as

$$(ff) = \frac{1 - [1 + A]c_\alpha}{1 - c_\alpha\left[\left(1 - \frac{1}{m_\alpha}\right) + \left(1 - \frac{1}{m_\beta}\right)A\right]}. \tag{S22}$$

Alternatively, the coverage with mode α in equation (S22) can be written as a function of the conditional probability $(ff)$ as,

$$c_\alpha = \frac{1 - (ff)}{1 + A - \left[1 - \frac{1}{m_\alpha} + \left(1 - \frac{1}{m_\beta}\right)A\right](ff)}. \tag{S23}$$

Then, the total coverage, $c = c_\alpha + c_\beta$ , is given by

$$c = \frac{1}{1 + \frac{\left(\frac{1}{m_\alpha} + \frac{A}{m_\beta}\right)}{1 + A}\frac{(ff)}{1 - (ff)}}. \tag{S24}$$

and the effective mode size $m_{eff}$ can be expressed, using equations (S18) and (S21), as

$$m_{eff} = \frac{c_\alpha m_\alpha + A c_\alpha m_\beta}{c_\alpha + A c_\alpha} = \frac{m_\alpha + A m_\beta}{1 + A}. \tag{S25}$$

The equations (S23), (S24), and (S25) give the partial coverage, total coverage, and effective coverage size in terms of model parameters and the conditional probability $(ff)$.

Now, to compute the equilibrium value of all these magnitudes $(A, c_\alpha, c_\beta, c, m_{eff})$, we need to know the conditional probability $(ff)$ as a function of the free ligand concentration $[SSB]$. To obtain this relation, we compute Eq. (S19) plus $m_\alpha/m_\beta$ times Eq. (S20), which simplifies to

$$0 = k_b^\alpha \frac{(ff)^{m_\alpha}}{1-(ff)}\left(c_\alpha + \frac{m_\alpha}{m_\beta}c_\beta\right) - k_u^\alpha c_\alpha, \tag{S26}$$

or equivalently,

$$\frac{k_u^\alpha}{k_b^\alpha} = \frac{(ff)^{m_\alpha}}{1-(ff)}\left(1 + \frac{m_\alpha}{m_\beta}\frac{c_\beta}{c_\alpha}\right). \tag{S27}$$

The latter equation, using the expressions for the binding constant, Eq. (S17), and for the coverage ratio, Eq. (S21), becomes

$$\frac{1}{\widehat{K_b^\alpha}\,[SSB]} = \frac{(ff)^{m_\alpha} + K_{\alpha\to\beta}(ff)^{m_\beta}}{1-(ff)}; \tag{S28}$$

which we restate as

$$[SSB] = \frac{1-(ff)}{\widehat{K_b^\alpha}\cdot\left((ff)^{m_\alpha} + K_{\alpha\to\beta}(ff)^{m_\beta}\right)}. \tag{S29}$$

This expression, Eq. (S29), is an implicit equation for the conditional probability $(ff)$ as a function of the free ligand concentration $[SSB]$.

In summary, for a given free ligand concentration $[SSB]$, we get the conditional probability $(ff)$ numerically solving Eq. (S29). The value of $(ff)$ then allows to compute the coverage ratio, $A \equiv c_\alpha/c_\beta$, using Eq. (S21), the coverage with mode $\alpha$, Eq. (S23), the total coverage, $c = c_\alpha + c_\beta = c_\alpha(1+A)$, and the effective mode size, $m_{eff}$, with Eq. (S25). Giving a set of parameters as in Table 1 of the main text, this procedure gives a prediction of these magnitudes as a function of the free ligand concentration $[SSB]$, as shown in Fig. 2 of the main text.

### 3. Fitting equilibrium predictions of the two-mode model to equilibrium experimental data of HmtSSB binding to ssDNA

We aim to fit the parameters of our two-mode model (described in the previous sections) to the experimental data in Fig. 3C and D of Ref. (Morin et al. 2017), which provide the effective mode, $m_{eff}$, and the total coverage, $c$, as a function of the free ligand concentration $[SSB]$. The equilibrium constants $\widehat{K_\alpha}$ and $K_{\alpha\to\beta}$ are the adjustable parameters. However, the size of the modes are set to $m_\alpha = 40$, and $m_\beta = 70$, as they were previously obtained from the experimental data in Ref. (Morin et al. 2017). [See Table 1 of Ref. (Morin et al. 2017).]

We determine the equilibrium constants by assuming a Gaussian likelihood and maximizing it, equivalent to minimizing

$$-\ln\mathcal{L} = \sum_{i=1}^{5}\frac{\left(c_i - c_{model}([SSB]_i)\right)^2}{\sigma_{ic}^2} + \sum_{i=1}^{5}\frac{\left(m_{eff_i} - m_{eff_{model}}([SSB]_i)\right)^2}{\sigma_{im_{eff}}^2}, \tag{S30}$$

where $c_i$, $m_{eff_i}$, and $[SSB]_i$ are the experimental values; $\sigma_{c,i}$ and $\sigma_{m_{eff},i}$ their experimental uncertainties; and $c_{model}$ and $m_{eff_{model}}$ the model predictions from expressions (S24) and

(S25), respectively. The minimization of the quantity in Eq. (S30) is a least squares minimization, but weighted with the uncertainty of each measurement.

The parameter values obtained from this fit are given in Table 1 of the main text, and the corresponding fitted curves are shown in Fig. 2 of the main text.

### 4. Fitting the kinetic model of SSB binding to ssDNA to the experimental data of the kinetics of HmtSSB and EcoSSB.

#### 4.1. Least squares fit

Fitting the kinetics has been revealed to be more challenging, requiring advanced fitting algorithms. For the kinetics fit, parameter estimation is performed via nonlinear least-squares using the Trust Region Reflective algorithm implemented in the `scipy.optimize.least_squares` function (Virtanen et al. 2020). This method minimizes the sum of squared residuals between model predictions and observed data. The Trust Region Reflective approach iteratively refines parameter estimates within a region around the current solution, adapting the step size based on local curvature and constraint boundaries (and using the same weight for all points in these fits).

The fit to the HmtSSB kinetic experimental data is shown in Fig. 3A of the main text, and the best-fit parameter values are in Table 2 of the main text. The fit is to the experimental data on the time evolution of the end-to-end extension of the ssDNA-HmtSSB complex during HmtSSB binding.

For the EcoSSB kinetics, the fit is shown in Fig. 4A of the main text, and the best-fit parameter values are reported in Table 3 of the main text (also shown here in Table S1, second column, left sub-column). In this case, the experimental data comprises the time evolution of the end-to-end extension, while EcoSSB is added to, removed from, and then added back to the solution.

#### 4.2. Bayesian analysis

In addition, we give a more refined study of the parameter space using Bayesian inference. We used the library PyMC (Salvatier, Wiecki, and Fonnesbeck 2016), with posterior sampling conducted via the No-U-Turn Sampler (NUTS) (Hoffman and Gelman 2014), a variant of Hamiltonian Monte Carlo. Convergence diagnostics and posterior summaries were computed using ArviZ (Kumar et al. 2019). The initial conditions are the least-squares fit values, shown in Table S1, second column, left subcolumn. Bayesian inference was performed using 16 independent chains with 5000 samples per chain; 2 chains were excluded due to poor convergence diagnostics. The posterior estimates for the model parameters were generally well-defined, with narrow credible intervals and low Monte Carlo standard errors, indicating precise estimation. A global good convergence between the chains was obtained, especially for the unbinding rate $k_u^{\alpha}$ and for the extension contraction in the $\beta$ mode, $\Delta x_{\beta}$. The values of the mean and the standard deviation of the parameters are given in Table S1, second column, right sub-column. In Figure S1, we show the pairwise joint posterior distributions of the model parameters. The diagonal panels display the marginal posterior distributions, while the off-diagonal panels show the joint distributions between each pair of parameters. Contours illustrate the density and potential correlations between parameters. The shape and orientation of these joint distributions provide insight into posterior dependency, i.e., correlations between parameters. For instance, as shown in Figure S1, the posterior distributions of several parameter pairs exhibit pronounced correlations, reflecting underlying degeneracies in the results of the fit to the experimental data. In particular, Fig. S1 shows a strong linear dependency between $\Delta x_{\beta}$ and $k_{\alpha\to\beta}$ suggesting a partial degeneracy: changes in one parameter can be compensated by reciprocal changes in the other while preserving the model fit. These degeneracies can be reduced with complementary experimental measurements, typically better of other magnitudes that have different dependencies on the model parameters.

#### 4.3. Kinetics for other concentrations of EcoSSB (extrapolation and least squares fits)

We obtained the previous results from a single EcoSSB concentration ($10nM$) with binding, unbinding, and rebinding experiments. Here, we ask to what extent the parameter values obtained are valid for other concentrations. The available experimental data provide only a partial answer, as for other concentrations, we have only binding data. In Panel A of Figure S2, we repeat (for comparison) the fit of the two-mode kinetic model shown in Fig. 4 of the main text. The parameter values of the fit are in Table 3 of the main text, and in Table S1 in the first column of numbers. In Panel B of Figure S2, we show how these fitted values extrapolate to predict the evolution of the extension of the EcoSSB-ssDNA complex at other EcoSSB concentrations. To extrapolate the binding rate, we assumed linear dependence on the SSB concentration, i.e.,

$$k_b^{\alpha}([SSB]) = k_b^{\alpha}([SSB]_0)\frac{[SSB]}{[SSB]_0}, \tag{S31}$$

in agreement with Eq. (S17). Panel B of Fig. S2 shows that this extrapolation is poor, providing only qualitative information and missing the quantitative prediction.

However, an overall fit to all data (see Panel C of Fig. S2) gives reasonable quantitative predictions across the range of concentrations for parameter values given in the second-to-last column of Table S1. A better fit is obtained by skipping the lower concentration ($0.1nM$) data, see Panel D of Fig. S2 for the fit and the last column in Table S1 for the parameter values. The parameter values of this last fit are more compatible with those obtained with the 10nM data alone (first two columns of numbers), except for the binding rate (evaluated at 10nM of EcoSSB). These results point toward the need for further investigation to understand the availability of the SSB in solution, due, for example, to its condensation in the free phase (Bonde et al. 2024).

| Parameter | *Bind - unbind 10 nM* | | All concentrations | All concentrations except 0.1 *nM* |
|---|---|---|---|---|
| | Least squares | Bayesian | | |
| $k_b^\alpha = k_b^8$ $(s^{-1})$ (at 10nM) | $9.1 \times 10^{-2}$ | $(9.1 \pm 0.4) \times 10^{-2}$ | $0.10$ | $2.4$ |
| $k_u^\alpha = k_u^8$ $(s^{-1})$ | $1.9 \times 10^{-2}$ | $(1.87 \pm 0.01) \times 10^{-2}$ | $1.4 \times 10^{-2}$ | $3.1 \times 10^{-2}$ |
| $k_{\alpha\to\beta} = k_{8\to17}$ $(s^{-1})$ | $0.41$ | $(0.41 \pm 0.03)$ | $1.4 \times 10^{-2}$ | $0.43$ |
| $k_{\beta\to\alpha} = k_{17\to8}$ $(s^{-1})$ | $5.3 \times 10^{-2}$ | $(5.3 \pm 0.1) \times 10^{-2}$ | $6.1 \times 10^{-2}$ | $4.0 \times 10^{-2}$ |
| $\Delta x_\alpha = \Delta x_8$ $(nm/nt)$ | $5.9 \times 10^{-2}$ | $(5.8 \pm 0.4) \times 10^{-2}$ | $-2.0 \times 10^{-2}$ | $-2.1 \times 10^{-2}$ |
| $\Delta x_\beta = \Delta x_{17}$ $(nm/nt)$ | $-0.22$ | $-(0.222 \pm 0.004)$ | $-1.1$ | $-0.18$ |

***Table S1: Parameter values of the two-mode binding model obtained for the EcoSSB kinetics*** *(at 12pN) experimental data from Ref.* (Naufer et al. 2021). *The values of the first two columns of numbers are for 10 nM of EcoSSB, obtained from a least squares fit (see Fig. 4 and Table 3 of the main text, and Fig. S2A) and from a Bayesian analysis (see Fig. S1). The last two columns are from least square fits, the second to last column for all concentrations show in Fig. S2C, and the last column for all concentrations of EcoSSB shown in Fig. S2D (i.e., skipping data for 0.1nM EcoSSB from the previous fit).*

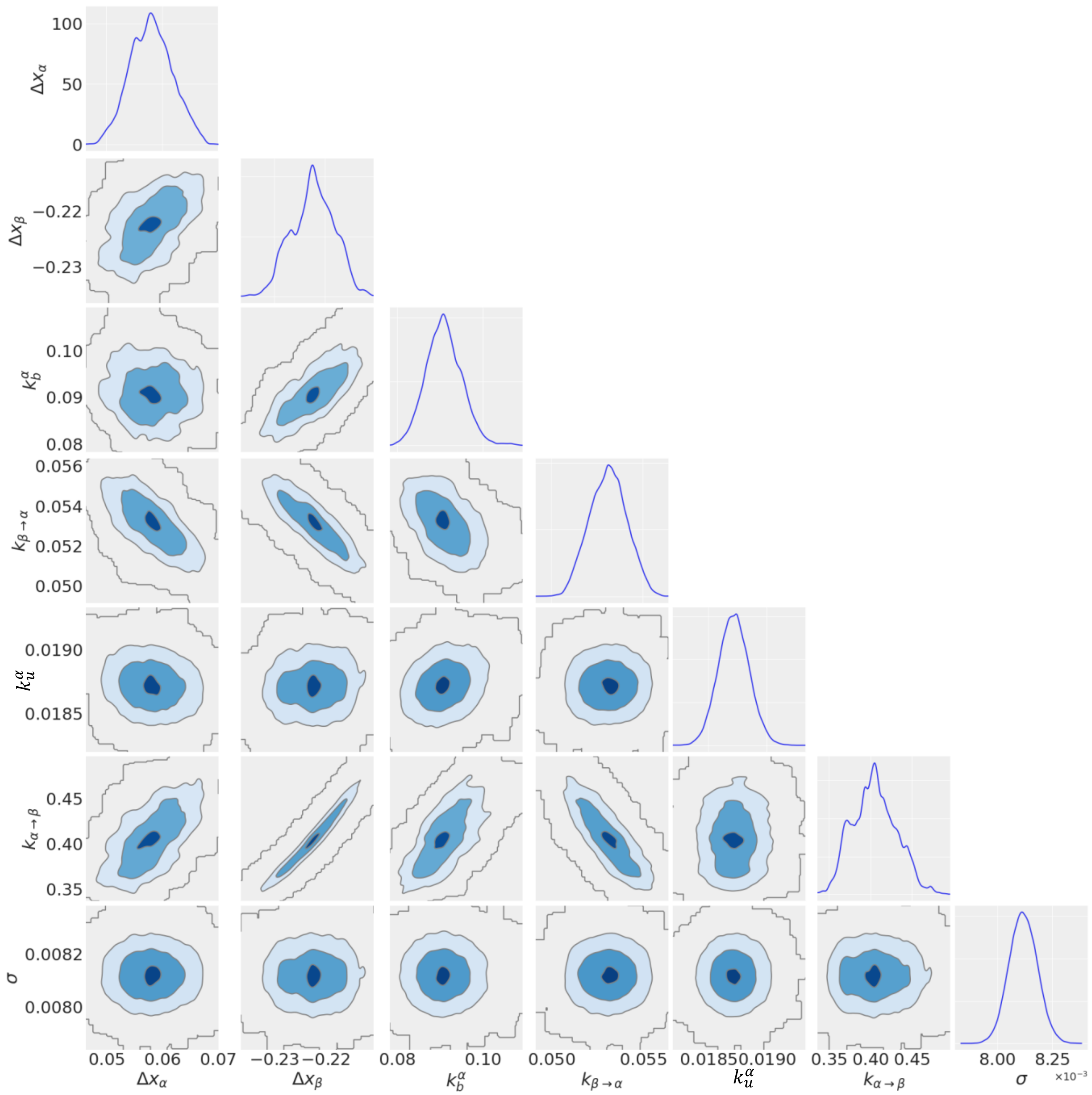


***Figure S1: Bayesian analysis of the two-mode kinetic model fitted to the EcoSSB experimental data of binding, unbinding and rebinding (at 10nM and 12pN) (Naufer et al. 2021).*** *Pair plot shows the marginal and joint posterior distributions for the inferred model parameters. Diagonal plots display marginal posteriors, while the off-diagonal plots show pairwise joint distributions with highest posterior density contours. Several parameter pairs display strong correlations (elongated joint posterior) indicating significant degeneracy between parameters.*

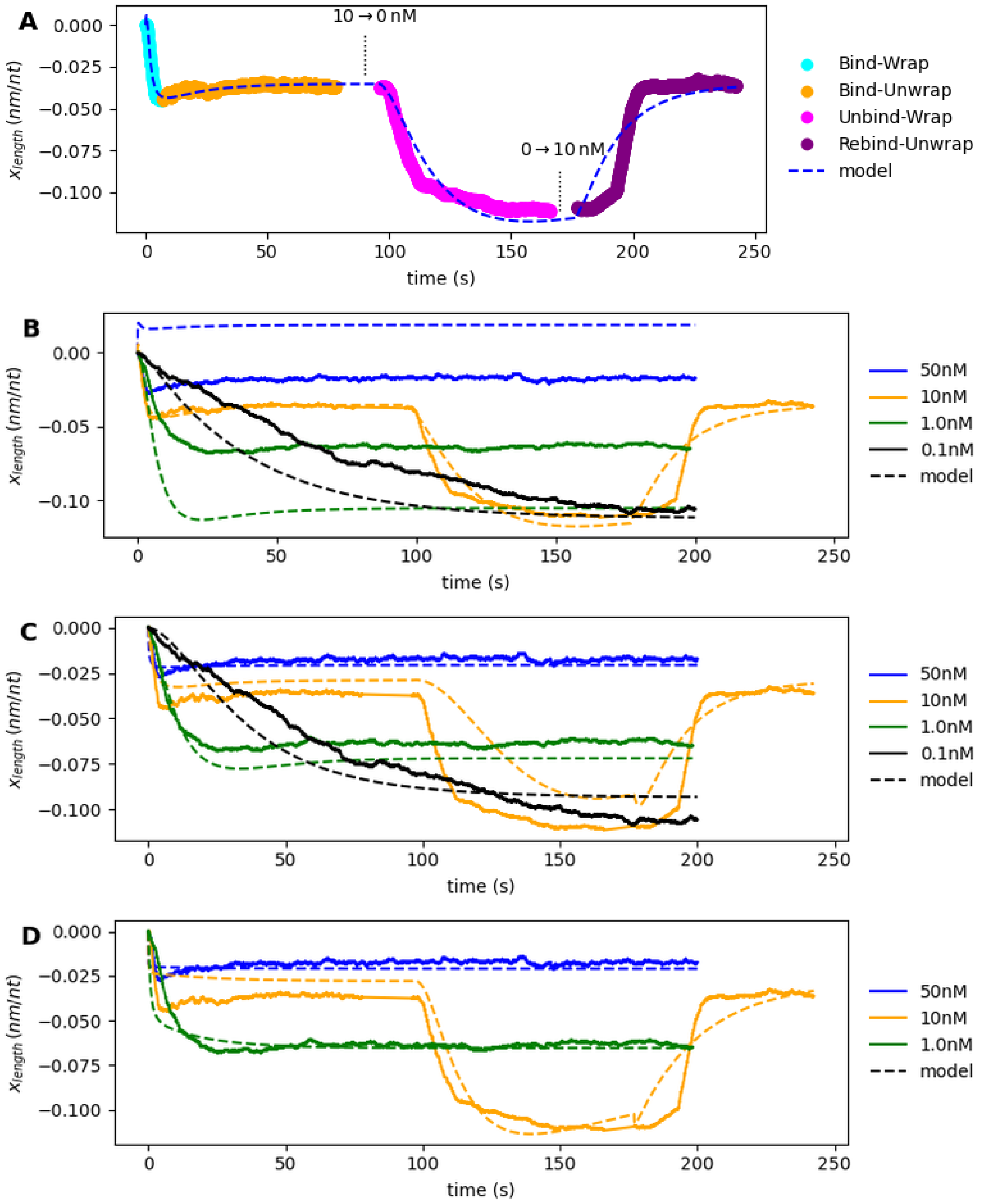


***Figure S2: Fits of the two-modes kinetic model to the experimental data of EcoSSB binding to ssDNA from Ref. (Naufer et al. 2021).*** *Panel A shows the fit of the two-modes model to the end-to-end experimental extension of the SSB-ssDNA complex as a function of time. The addition, elimination, and readdition of EcoSSB gives rise to a binding, unbinding, and rebinding kinetics, whose implications for the end-to-end extensions of the SSB-ssDNA complex evolution is well explained by the two-modes kinetic model. (This kinetics underlying this case is further analysed in Fig. 4 of the main text. The parameter values of the fit are given in Table 3 of the main text and in the first column of numbers in Table S1.) Panel B show the extrapolations for other concentrations [assuming linearity of the binding rate with the EcoSSB concentration, Eq. (S31)]. Panel C shows the same experimental data as in panel B, but with the best fit using all the data. Panel D skips the 0.1nM from the representation and the fit. (The parameter values of these two fits are shown in the two last columns of Table S1).*

Suksombat, Sukrit, Rustem Khafizov, Alexander G Kozlov, Timothy M Lohman, and Yann R Chemla. 2015. “Structural Dynamics of *E. Coli* Single-Stranded DNA Binding Protein Reveal

DNA Wrapping and Unwrapping Pathways.” *ELife* 4: 1–23. https://doi.org/10.7554/eLife.08193.

Villaluenga, Juan P. G., Jules Vidal, and Francisco Javier Cao-García. 2020. “Noncooperative Thermodynamics and Kinetic Models of Ligand Binding to Polymers: Connecting McGhee–von Hippel Model with the Tonks Gas Model.” *Physical Review E* 102 (1): 012407. https://doi.org/10.1103/PhysRevE.102.012407.

Villaluenga, Juan P.G., and Francisco Javier Cao-García. 2022. “Cooperative Kinetics of Ligand Binding to Linear Polymers.” *Computational and Structural Biotechnology Journal* 20: 521–33. https://doi.org/10.1016/j.csbj.2021.12.043.

Villaluenga, Juan P G, David Brunete, and Francisco Javier Cao-García. 2023. “Competitive Ligand Binding Kinetics to Linear Polymers.” *Physical Review E* 107 (2): 024401. https://doi.org/10.1103/PhysRevE.107.024401.

Virtanen, Pauli, Ralf Gommers, Travis E. Oliphant, Matt Haberland, Tyler Reddy, David Cournapeau, Evgeni Burovski, et al. 2020. “SciPy 1.0: Fundamental Algorithms for Scientific Computing in Python.” *Nature Methods 2020 17:3* 17 (3): 261–72. https://doi.org/10.1038/s41592-019-0686-2.
